# Development of quantitative high-throughput screening methods for identification of antifungal biocontrol strains

**DOI:** 10.1101/2021.06.23.449687

**Authors:** Bodil Kjeldgaard, Ana Rute Neves, César Fonseca, Ákos T. Kovács, Patricia Domínguez- Cuevas

## Abstract

Large screens of bacterial strain collections to identify potential biocontrol agents are often time consuming, costly, and fail to provide quantitative results. In this study, we present two quantitative and high-throughput methods to assess the inhibitory capacity of bacterial biocontrol candidates against fungal phytopathogens. One method measures the inhibitory effect of bacterial culture supernatant components on the fungal growth, while the other accounts for direct interaction between growing bacteria and the fungus by co-cultivating the two organisms. The antagonistic supernatant method quantifies the culture components’ antifungal activity by calculating the cumulative impact of supernatant addition relative to a non-treated fungal control, while the antagonistic co-cultivation method identifies the minimal bacterial cell concentration required to inhibit fungal growth by co-inoculating fungal spores with bacterial culture dilution series. Thereby, both methods provide quantitative measures of biocontrol efficiency and allow prominent fungal inhibitors to be distinguished from less effective strains. The combination of the two methods shed light on the type of inhibition mechanisms and provide the basis for further mode of action studies. We demonstrate the efficacy of the methods using *Bacillus spp*. with different levels of antifungal activities as model antagonists and quantify their inhibitory potency against classic plant pathogens.

**Importance:** Fungal phytopathogens are responsible for tremendous agricultural losses on annual basis. While microbial biocontrol agents represent a promising solution to the problem, there is a growing need for high-throughput methods to evaluate and quantify inhibitory properties of new potential biocontrol agents for agricultural application. In this study, we present two high-throughput and quantitative fungal inhibition methods that are suitable for commercial biocontrol screening.

## Introduction

On annual basis, it is estimated that the global crop production suffers losses between 20 to 40 percent due to pests and plant diseases [1]. Plant diseases alone are predicted to cost the global economy a staggering $220 billion per year [2]. Among other plant pathogens, fungal phytopathogens contribute to considerable losses in agriculture and greatly impact food security in developing countries [3–5]. Not only do fungal pathogens affect the yield, but fungal crop infections also lead to severe reductions of post-harvest crop quality. For instance, accumulation of high levels of mycotoxins render crops unsafe for human consumption and for animal forage [6,7]. In modern intensified agriculture, fungal diseases are commonly fought using fungicides [8], but the rising fungicide resistance and chemical pollution represent a challenge to the sustainable use of these chemicals in agriculture [9–12]. In addition, many fungicides are hazardous to humans and infer developmental or reproductive toxicity [13]. Application of microbial biocontrol agents represent a safe alternative to the intensive use of agrochemicals [11,14]. Biocontrol agents derived from plant growth promoting rhizobacteria (PGPR) reside in close association with plant roots, and protect the plant from phytopathogens by priming the plant defense response, competing for nutrients and/or directly antagonizing growth and development of the pathogenic intruders [15,16]. Strains from *Pseudomonas, Burkholderia, Streptomyces* and the *Bacillus* genera are well known for their antifungal capacity and for the production of a large variety of bioactive metabolites [14,17–22]. Although, the inhibitory effect of specific soil bacteria is well documented and recognized, there is a lack of quantitative and HTP screening procedures to identify competent biocontrol agents. Consequently, many potential biocontrol agents eventually fail to suppress plant diseases in field trials [23,24]. Classic antagonistic screens assess the impact of the biocontrol candidates on the phytopathogen after co-inoculation on solid media, which are referred to as dual-culture, plate confrontation or inhibition zone assays [23,25]. Such methods account for numerous factors including nutrients or space competition, cell surface components and the induced or constitutive secretion of volatile or soluble metabolites [19,25–27]. Other antagonistic assays evaluate the effect of individual inhibitory components on the phytopathogens’ growth like volatiles, polyketides, lipopeptides, siderophores and lytic enzymes including, chitinases, glucanases, and proteases [19,25]. More complex antagonistic assays, such as leaf disc or seedlings assays [28,29], investigate the tripartite interaction between biocontrol candidate, phytopathogen and plant host, while non-antagonistic assays assess the importance of complementary inhibitory mechanisms including niche colonization and priming of the plant immune response [25]. Nevertheless, most screening systems are low throughput and provide only semi-quantitative measurements of the inhibition potential against the fungus. Therefore, there is a need to develop more efficient screening methods combining quantitative measurements of antimicrobial activity with automation to increase speed and reduce resources required for the identification of good candidates.

Here, we describe two fungal inhibition methods to evaluate antifungal potency of potential biocontrol agents. Both methods accommodate high-throughput (HTP) screening of bacterial biocontrol candidates and provide accurate quantification of their inhibitory capacity. The major difference between the two methods is represented by the use of growing bacterial cells as opposed to (cell-inactive) culture supernatants. We demonstrate the efficacy of the methods using bacterial strains with different antifungal performance. Using the two novel methods, antifungal properties of bacteria were compared, and prominent fungal inhibitors were distinguished from less effective bacterial strains. Both methods were developed utilizing *Fusarium culmorum* as model plant pathogen and *Bacillus* spp. as model antagonists. In addition, the methods were further applied for screening *Bacillus spp.* against other important phytopathogens, i.e. *Fusarium graminearum* and *Botrytis cinerea*, proving that our method can readily be adjusted to other fungal species.

## Results

### Co-inoculation of fungal spores with bacterial dilution series facilitates quantification of inhibition potency

The so-called dual-culture assay is among the most common screening methods to identify potent fungal inhibitors from microbial collections [17,21,30–32]. Typically, the assay is performed by inoculating potential biocontrol agents at a fixed distance from the pathogenic fungal inoculum on a petri dish as illustrated in Fig. 1A. Subsequently, the biocontrol agent’s ability to suppress fungal growth is manually assessed by measuring the radius of the mycelial growth relative to the control or the size of the inhibition zone [33–36]. However, accurate comparison and subsequent ranking of large numbers of strains is difficult with this assay due to the format of the readout. To improve the evaluation and accurate quantification of antifungal potency, we developed a HTP fungal inhibition assay based on direct co-inoculation of bacterial cultures and fungal plant pathogenic spores. First, bacterial overnight cultures were normalized to the same optical density at 600 nm (OD_600_) and 5-fold serial dilutions were prepared (down to an amount of approximately 3-5 bacterial cells inoculated). The OD_600_ of each strain used in the study was correlated to viable cell counts (colony forming units (CFUs)). A fungal spore suspension was prepared and mixed thoroughly to ensure a homogeneous spore distribution. Then, the bacterial dilution series were co-inoculated with a fixed quantity of fungal spores to determine the minimal inhibitory cell concentration (MICC) that abolishes fungal growth (Fig. 1B). With this setup, a low MICC indicates a higher antifungal potency for a given bacterial strain. The assay was prepared on appropriate agar medium for fungal cultivation in 48-well microtiter plates and incubated 5 days at room temperature before assessing the fungal growth.

**Fig. 1:**
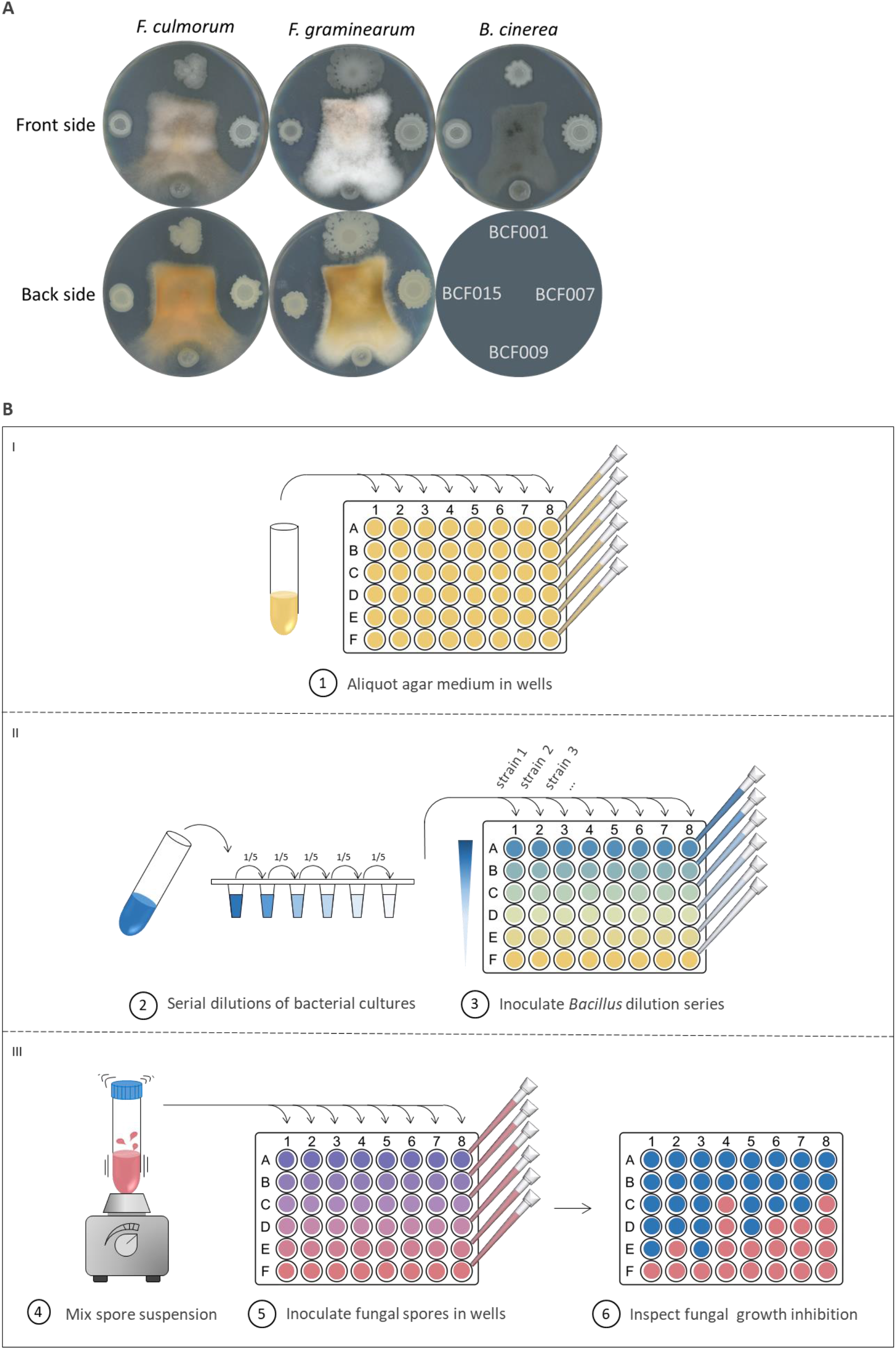
Fungal inhibition assays. (A) *B. subtilis* BCF001, *B. amyloliquefaciens* BCF007, *B. paralicheniformis* BCF009 and *B. velezensis* BCF015 were spotted around central inocula of *F. culmorum*, *F. graminearum* and *B. cinerea* on agar medium. Inhibition zones were observed 4-5 days post inoculation. (B) In each well of a 48-well microtiter plate, molten PDA medium was aliquoted and let to solidify. 5-fold dilution series of *Bacillus* strains were prepared from cultures normalized to the same OD_600_. In each column, consecutive wells were co-inoculated with the *Bacillus* dilution series and a fixed quantity of fungal spores (constant volume of fungal spore suspension) on the agar surface. The spore suspensions were mixed thoroughly before aliquoting. Assay results were evaluated by visual inspection after 5 days incubation at room temperature.

Four *Bacillus* strains of different species were selected based on their differential properties for fungal inhibition. The inhibitory capacity of the selected strains, *Bacillus subtilis* BCF001, *Bacillus amyloliquefaciens* BCF007, *Bacillus paralicheniformis* BCF009 and *Bacillus velezensis* BCF015, was quantified against *F. culmorum* DSM1094 using the developed method (Fig. 2B). The assay was conducted as three biological independent replicates, rendering very similar results, which indicates that the method is highly reproducible.

**Fig. 2:**
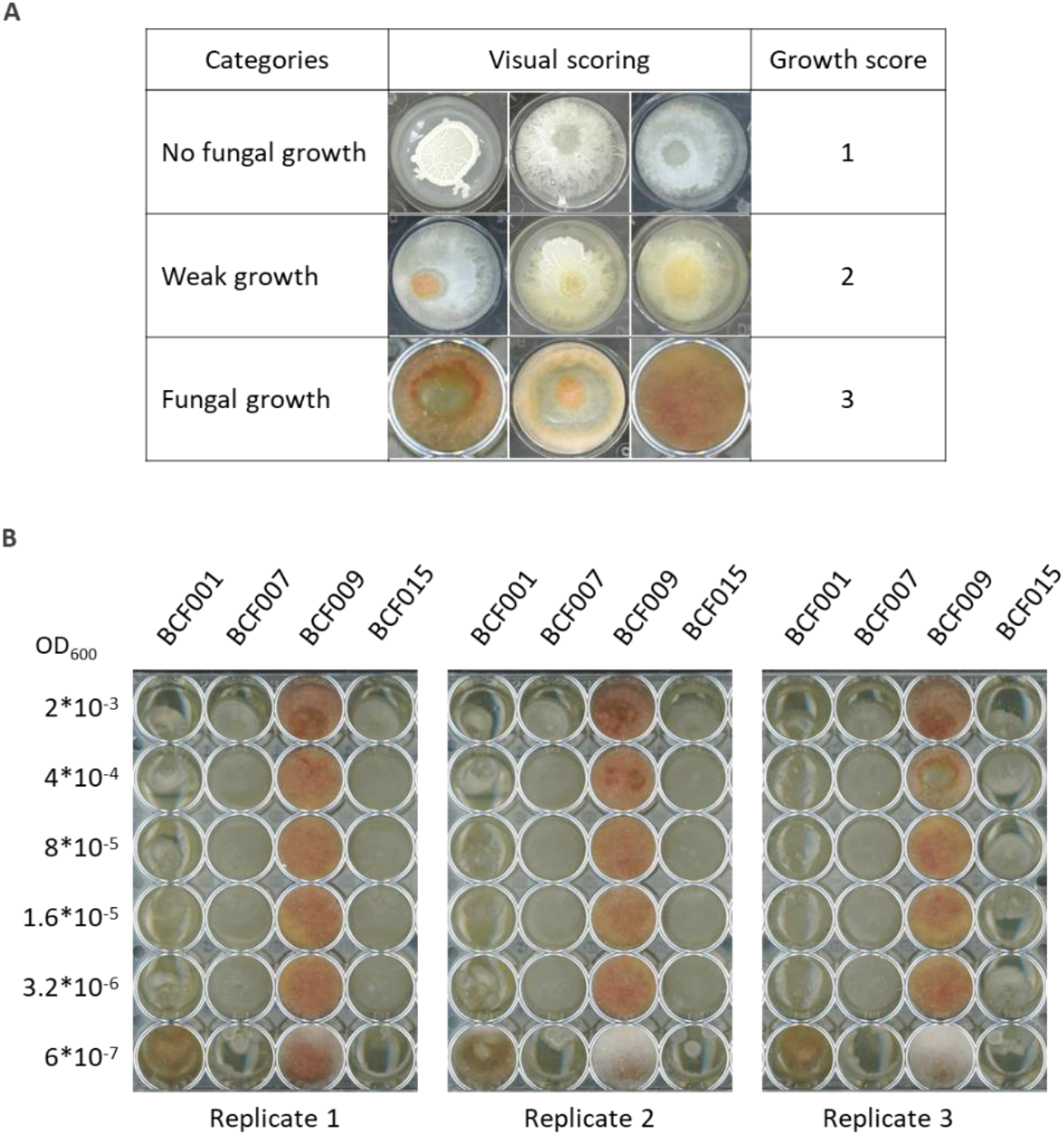
Comparison of *Bacillus* spp. inhibitory properties against *F. culmorum*. (A) *F. culmorum* growth was scored following a three steps scale illustrated with three image examples per category. (B) Dilution series of *B. subtilis* BCF001, *B. amyloliquefaciens* BCF007, *B. paralicheniformis* BCF009 and *B. velezensis* BCF015 were prepared and inoculated in consecutive columns of a 48-well microtiter plate. A constant spore concentration of *F. culmorum* was inoculated in each well. ODs of serial dilutions are indicated on the left panel (OD_600_).

Fungal growth inhibition was scored as *no growth (1)*, *weak growth (2)* and *(positive) growth (3)* (Fig. 2B). Fungal growth was defined by presence of visible hyphae in the well, including those that where half-covered by fungal mycelia. No fungal growth was defined by complete absence of visible fungal mycelia. Weak growth was defined by the presence of barely visible hyphae in all replicates, or by the absence of growth in half of the replicates. The weak growth category included the wells where *F. culmorum* produced (orange) pigmentation, even in the absence of visible hyphae.

Comparison of bacterial MICC allowed ranking of the strains in accordance to their fungal inhibition properties (Fig. 3A, Table S1). The strains *B. amyloliquefaciens* BCF007 and *B. velezensis* BCF015 showed the most potent inhibition properties, and even at the highest dilution step (corresponding to an initial OD_600_ of 6.4*10^−6^, equivalent to 3-5 CFUs inoculated), the two strains were able to inhibit *F. culmorum* growth. Determination of the minimal cell concentration that inhibits fungal growth even more accurately would require to conduct the assay using serial dilutions at smaller dilution factors. Nevertheless, the experimental setup is optimized for HTP screening of large strain collections. *B. subtilis* BCF001 also displayed potent fungal inhibition properties, and abolished fungal growth up to an OD_600_ of 3.2*10^−5^, corresponding to 19 CFU inoculated. *B. paralicheniformis* BCF009, however, did not affect fungal growth even at the lowest dilution step, corresponding to an OD_600_ of 4*10^−3^, 3.89*10^4^ CFUs inoculated.

**Fig. 3:**
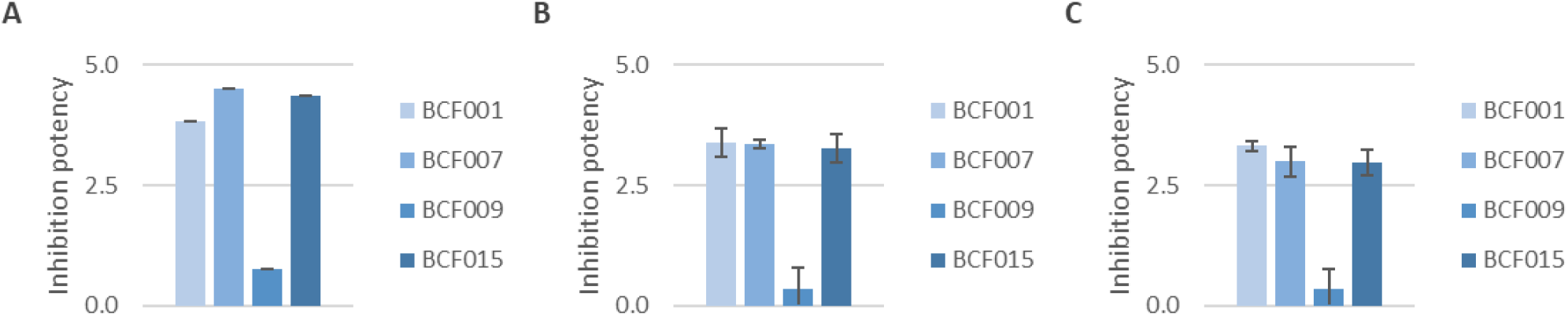
Inhibition potency of *Bacilli* against fungal phytopathogens. The inhibition potency of *B. subtilis* BCF001, *B. amyloliquefaciens* BCF007, *B. paralicheniformis* BCF009 and *B. velezensis* BCF015 was assigned a numerical score based on the minimal inhibitory cell concentration (MICC) against (A) *F. culmorum*, (B) *F. graminearum* and (C) *B. cinerea.* Inhibition scores were calculated by applying the formula #1.

To generate visual and directly comparable plots of the inhibition results, we calculated a numerical inhibition score that reflects the inhibitory capacity of each strain. In brief, the lowest cell concentration that caused fungal growth inhibition was identified for each strain, and a numerical inhibition score was calculated based on the natural logarithm to the MICC, by applying the empirical formula #1 (see material and methods section) (Fig. 3A, Table S1). Plotting the inhibition scores clearly indicated that *B. amyloliquefaciens* BCF007 and *B. velezensis* BCF015 were the most efficient strains inhibiting growth of *F. culmorum*. *B. subtilis* BCF001 also showed a high inhibition score although smaller than the ones calculated for the two former strains. Scoring results for *B. paralicheniformis* BCF009 matched the low inhibitory activity of this strain against *F. culmorum*.

### Evaluation of inhibition method with additional fungal phytopathogens

Using the newly developed method, the antifungal activity of *B. subtilis* BCF001, *B. amyloliquefaciens* BCF007, *B. paralicheniformis* BCF009 and *B. velezensis* BCF015 was also quantified against other phytopathogenic filamentous fungi, specifically against *F. graminearum* and *B. cinerea* strains. Images acquired to record the experimental results, the calculated MICCs and inhibition scores can be found in Fig. S1, Table S1. Interestingly, our inhibition assay proved applicable to these additional species of plant pathogenic fungi. The initial spore concentration was the only parameter that required adjustment when testing growth inhibition against the new fungal strains.

The previously described scoring system was applied to the results obtained for the three fungal species (Fig. 3). While *B. amyloliquefaciens* BCF007 and *B. velezensis* BCF015 displayed the most efficient growth inhibition of *F. culmorum*, differences between them and *B. subtilis* BCF001 were minimal when assayed against *F. graminearum* DSM4528. Furthermore, the strain *B. subtilis* BCF001 proved more efficient against *B. cinerea* Kern B2 compared to *B. amyloliquefaciens* BCF007 and *B. velezensis* BCF015. Inhibition scoring results obtained for *B. paralicheniformis* BCF009 were in accordance with the poor inhibitory properties of this strain, regardless of the fungal pathogen tested.

### Inoculation of fungal spores with bacterial culture supernatants validates the inhibitory importance of supernatant components

While the antagonistic co-inoculation method assesses several inhibitory mechanisms, the use of cell-inactive supernatants accounts for the bioactivity of secreted metabolites, like enzymes, lipopeptides and polyketides, on the fungus. In the antagonistic supernatant method, fungal growth was estimated by OD_600_ measurements in liquid cultures and therefore, reduced fungal growth in the presence of bacterial metabolites reflects greater inhibition potency of a given strain. For this method, bacterial cultures were grown overnight and normalized to the same OD_600_ value prior to collection of the culture supernatants by centrifugation (Fig. 4). A range of increasing supernatant volumes (10-80 μl) were inoculated into fungal spore suspensions prepared in liquid medium in 48-well microtiter plates. To avoid any possibility of misreads arising from remaining bacterial cells present in the supernatants, bacteriostatic antibiotics were added to the fungal spore suspension to prevent bacterial growth. The microtiter plates were incubated statically at 25°C in darkness. Subsequently, the fungal growth was quantified by spectrophotometric measurement.

**Fig. 4.**
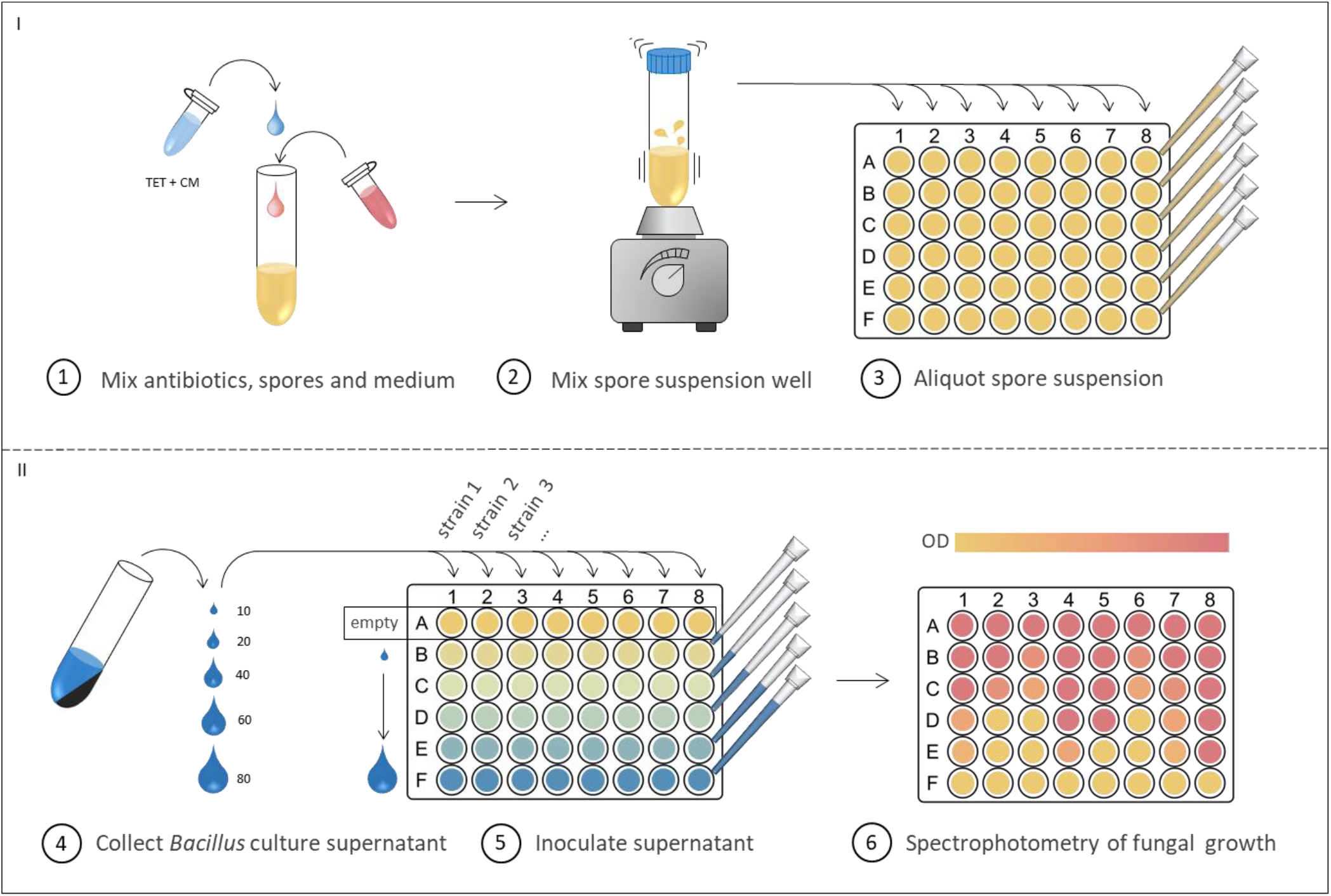
Supernatant inhibition method. A fungal spore suspension was prepared with PDB medium and the bacteriostatics tetracycline (TET) and chloramphenicol (CM). The suspension was mixed well and aliquoted into each well of a 48-well microtiter plate. *Bacillus* overnight cultures were adjusted to 2 OD_600_ and were subsequently centrifuged to collect the supernatants. In each column, *Bacillus* supernatant volumes (10-80 μl) were added to consecutive wells. Fungal growth was evaluated by spectrophotometric measurements after 5 days incubation at 25°C in darkness.

The inhibition assay demonstrated a reverse correlation between the added volume of bacterial cell-inactive supernatant and growth of *F. culmorum*, *F. graminearum* or *B. cinerea* (Fig. 5A-C). For potent fungal inhibitory strains, including *B. subtilis* BCF001, *B. amyloliquefaciens* BCF007 and *B. velezensis* BCF015, increasing volumes of bacterial supernatant correlated with a progressive decrease of fungal growth. Addition of *B. paralicheniformis* BCF009 supernatant impacted fungal growth to a much lesser extent, which is in accordance with the results obtained with the co-inoculation method. In some cases, a slight increase in fungal growth was observed in response to addition of the largest supernatant volume (80 μl culture supernatant), suggesting an effect of supplemented nutrients (not consumed by the bacteria), which outweighed the antifungal bioactive metabolites.

**Fig. 5:**
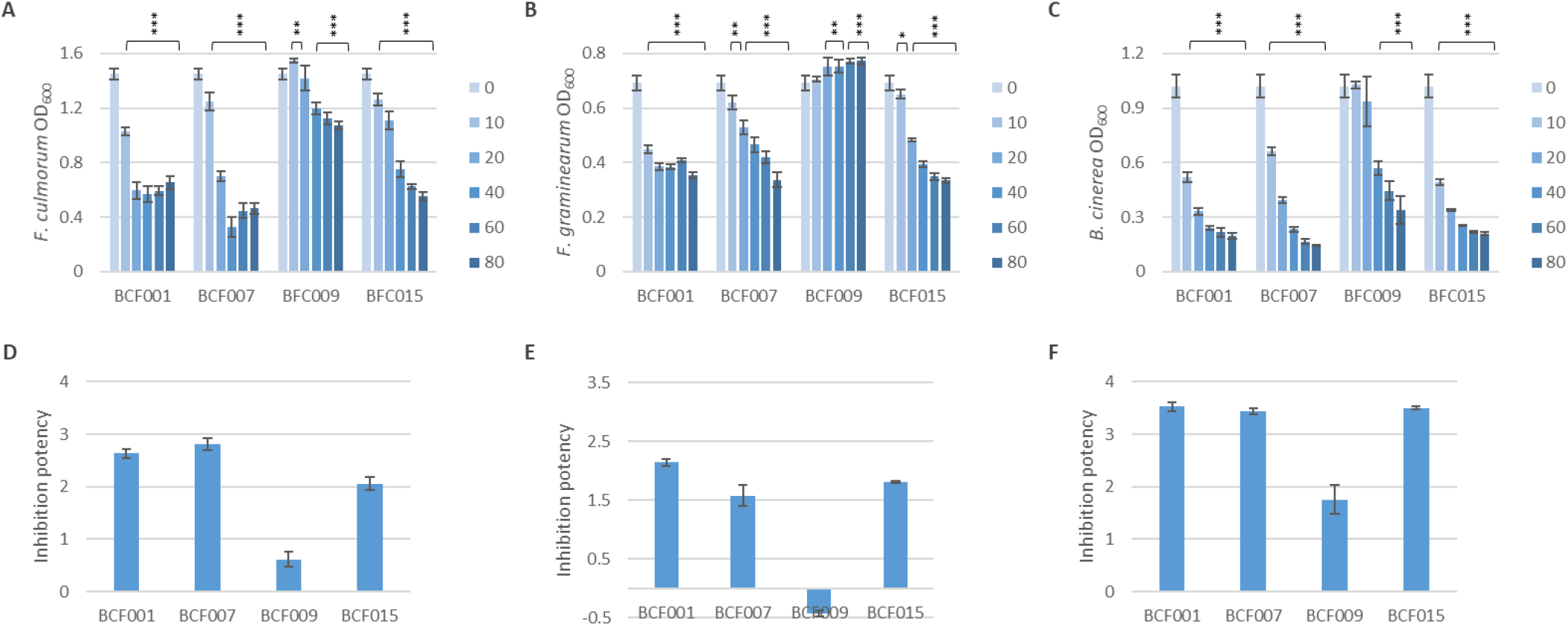
Fungal Growth inhibition by *Bacillus spp.* culture supernatants. *Bacillus spp.* cultures were grown overnight in LB broth and normalized to OD_600_ 2. Different volumes of cell-inactive supernatants (10, 20, 40, 60 and 80 μl) were added to 48-well microtiter plates containing PDB medium and a fixed fungal spore concentration. Bacterial growth was inhibited by presence of bacteriostatic antibiotics in the culture medium (50 μg/ml chloramphenicol and tetracycline 10 μg/ml, respectively). Plates were incubated at 25°C without shaking for 60 h (in darkness) and fungal growth was measured by spectrophotometry at 600 nm. Plots correspond to inhibition results for (A) *F. culmorum*, (B) *F. graminearum* and (C) *B. cinerea*. Statistical significance for each supernatant volume compared to the control was calculated based on biological triplicates. *P<0.05, **P<0.005, ***P<0.0005. For quantification of the culture supernatant inhibition potency, an inhibition score was assigned to each *Bacillus* strain by subtracting the accumulated relative fungal growth of (D) *F. culmorum*, (E) *F. graminearum* and (F) *B. cinerea* in response to all supernatant volumes from the total potential growth according to Equation #2. Standard deviations were calculated based on biological triplicates.

### Scoring of supernatant inhibition potency

A numerical inhibition score was assigned to each strain by quantifying the fungal growth in response to supernatant addition (5 volumes). The inhibition scores were calculated by subtracting the accumulated relative fungal growth from total potential fungal growth in the 5 conditions tested (Equation #2, see material and methods section).

The inhibition scores can be found in Table S1. Culture supernatants of *B. subtilis* BCF001 and *B. amyloliquefaciens* BCF007 showed the most effective growth inhibition of *F. culmorum* (Fig. 5A and D). Against *F. graminearum*, *B. subtilis* BCF001 culture supernatants proved most inhibitory, while *B. subtilis* BCF001, *B. amyloliquefaciens* BCF007 and *B. velezensis* BCF015 all displayed similar inhibition potency of *B. cinerea*. *B. paralicheniformis* BCF009 culture supernatants poorly inhibited *F. culmorum* growth.

It is worth mentioning that assays prepared from different fungal spore stock solutions showed small variations in final fungal growth (Fig. S2), possibly due to variations in the initial spore count or spore viability. These variations could be minimized either by prolonging the incubation time until readout or by preparing assays from a unique spore stock. For this reason, biological replicates were prepared with the same fungal spore solution. Nevertheless, the overall conclusions remained unchanged despite variations in the final fungal growth.

### Comparison of methods to remove or inactivate cells from culture supernatants

Typically, bacterial cell-inactive supernatants are generated by filtration of the cell cultures [37–40]. In contrast, in the proposed setup cell-inactive supernatants were generated by centrifugation to pellet bacterial cells. To avoid cell growth of remaining bacteria in suspension, we added the bacteriostatic antibiotics chloramphenicol and tetracycline. Plating of the fungal growth suspension on bacteria-selective medium after the end point measurement produced no bacterial growth, indicating that the antibiotics effectively inhibited growth of any putative remaining bacterial cells (data not shown). Final fungal growth measurements reached by *F. culmorum*, *F. graminearum* and *B. cinerea* were somewhat affected by addition of the respective antibiotics (Fig. S3). The results obtained with filtered supernatants with and without antibiotics revealed that the relative inhibition results remained unaltered, although the addition of antibiotics slightly but significantly reduced the fungal growth (Fig. S4, Table S2). Both filtration and antibiotic addition had generally no effect on the fungal inhibitory potency, suggesting that bioactive metabolites remain active after both procedures (Fig. S5, Table S2).

### Comparison of the two methods proposed in the present study and the dual-culture assay

The two proposed quantitative HTP methods were compared to the common dual-culture assay using plates inoculated with fungus and *Bacilli* (Fig. 1A). In accordance with results from the two methods, zones of fungal growth inhibition were observed around the bacterial colonies of *B. subtilis* BCF001, *B. amyloliquefaciens* BCF007 and *B. velezensis* BCF0015, whereas growth of *F. culmorum*, *F. graminearum* and *B. cinerea* was nearly unaffected by *B. paralicheniformis* BCF009. Although, the traditional dual-culture assay allowed assessment of fungal inhibitory capacity, the results lack the accurate quantification of the bacterial inhibition potency provided by the proposed methods. In addition, the simple numerical scoring systems allow easy comparison between a large number of strains.

While the antagonistic supernatant method specifically evaluates the inhibition potency of secreted metabolites, like enzymes, lipopeptides and polyketides, the antagonistic co-inoculation method quantifies the inhibitory effect of actively growing bacteria, and thereby accounts for additional factors like competition for nutrients and space. Therefore, differences in results obtained by the two methods could provide insights on the mode of action and serve as the starting point for in-depth characterization of their molecular inhibitory mechanism.

## Discussion

Despite the steadily growing market share of biocontrol products compared to the use of conventional pesticides [15,41], plant pathogen management continues to heavily rely on chemical substances with associated risks to human health and the environment [42]. Regardless of the increased (research) efforts on microbial biocontrol product development, their deployment into market is difficult due to the absence of a global harmonized framework, public misinformation, complex regulation and registration procedures as well as expensive field trials [43–46]. The application of reliable methods to identify and select potent biocontrol strains is of critical importance to reduce product development costs and accelerate their market implementation.

Large screens of strain collections are often laborious and expensive, where the complexity of the screening system is proportional to the cost [47]. Simple HTP screens are cost-effective, however they often fail to provide quantitative results for accurate comparison of the biocontrol strains. Furthermore, following the HTP screening and candidate selection, many strains eventually show low efficacy in field trials, demonstrating a discrepancy between *in vitro* laboratory conditions and *in planta* application experiments. To reduce costs and minimize failed tests, it is therefore crucial to identify biocontrol candidates in primary screens before moving on to complex experimental systems such as greenhouses or field trials. Accordingly, development of simple, reliable and quantitative HTP screening methods is crucial for ranking and selection of the best biocontrol candidates.

In this study, bacterial strains were ranked according to their bioactivity against fungal plant pathogens using two novel HTP antagonistic methods: i) an antagonistic co-cultivation method based on the inoculation of a constant number of fungal spores with bacterial dilutions series on a solid media to generate a quantitative measurement of the minimal inhibitory (bacterial) cell concentration (MICC) of fungal growth and, ii) an antagonistic cell-inactive supernatant method based on the inoculation of a constant number of fungal spores with different volumes of bacterial culture supernatants in a liquid media to provide a quantitative measurement of fungal growth inhibition by secreted metabolites.

Although classic antagonistic methods that assess adjacent growth of two species on agar plates allow assessment of fungal inhibition potential [17,30–32,48], their accuracy is limited. Factors such as inoculum size, diffusion rate of metabolites, and differential conditions between agar plates contribute to the inaccuracy of classic antagonistic assays and impairs ranking of biocontrol candidates. The quantification provided by the proposed antagonistic co-cultivation method allowed ranking of biocontrol candidates according to their inhibitory strength by means of MICC values. In addition, the direct co-inoculation of fungal spores with bacterial cells do not only assesses the impact on mycelial growth, but also on fungal spore germination, contrary to the classic agar plate methods. These type of antagonistic methods, employing direct co-inoculation of biocontrol candidate and pathogen have been developed for other applications, for instance lactic acid biocontrol bacteria co-inoculated with food spoilage fungi in dairy industry [49,50]. However, these assays lack precise quantification of inhibition potency, regardless of the similarities with the co-inoculation method described here.

Compared to simple dual-culture assays like ours, the more complex *in planta* assays include the tripartite interaction of biocontrol candidate, phytopathogen and plant host [28,29,51], and thereby attempt to mimic field settings. Despite the closer resemblance to natural conditions, the complexity of these screening systems limits the throughput (Haidar et al., 2016) and leads to additional variation [52], which complicates the interpretation of results compared to our method. However, screens that consider the interaction between three biological systems may be advantageous to apply to a subset of biocontrol candidates following an initial HTP screen [53,54].

While the antagonistic co-cultivation methods investigate the direct interaction between pathogen and biocontrol candidate, the supernatant antagonistic methods estimate the inhibition potency of secreted bioactive metabolites from the biocontrol candidate on the pathogen. The experimental setup is either agar- or liquid-based, like the proposed antagonistic cell-inactive supernatant method described in this study. Agar-based screens rely on evaluation of fungal colony growth or inhibition zones in response to addition of supernatant from the biocontrol candidates [32,38,55]. Some assays allow a quantitative comparison of inhibition capacity between strains by calculating the reciprocal to the highest supernatant dilution that exhibits a clear zone of inhibition [38,55]. Even so, the scoring is notably laborious and low-throughput compared to the proposed antagonistic supernatant method described in this study.

Liquid-based supernatant assays depend on the assessment of fungal morphology by microscopy [55,56], evaluation of fungal growth by dry weight [57], or spectrophotometry in response to the biocontrol supernatants [34,37]. The latter share great resemblance to the proposed cell-inactive supernatant method and allows HTP and reproducible assessment of the antifungal metabolite potency. However, the accurate quantification of antifungal capacity by determination of supernatant inhibition scores in the proposed method allows easy benchmarking and comparison between biocontrol supernatants, which represents a major advantage compared to previously reported methods.

Both assays were initially developed utilizing *F. culmorum* as model plant pathogen, and different *Bacillus* species as model antagonists. Subsequently, the assays were validated with the relevant plant pathogenic fungi *B. cinerea* and *F. graminearum*. The results demonstrated that the methods are robust and can readily be adapted for different fungal species. Moreover, the methods can be adapted to assess the potential for inhibiting bacterial plant pathogens. In agriculture, not only fungal plant pathogens, but also bacterial plant pathogens contribute to significant yield losses [58]. The adaptation of the proposed methods to quantify inhibition potency against plant pathogenic bacteria simply requires a fluorophore-labelled bacterial pathogen, a bacterial pathogen with a selective marker (i.e. antibiotic resistance) or a pathogen-selective growth medium in order to make it applicable for HTP screening. For discrimination between bacterial strains with similar inhibition potency, the antagonistic co-cultivation method can be adjusted by using a smaller dilution factor. Furthermore, the dilution factors can be adjusted to fit more or less potent biocontrol candidates. Finally, the applicability of the methods may even be extended to other areas such as biocontrol screens against human or animal pathogens or screens against food spoilage microorganisms.

Several previous studies employ combinatory screens with two or more assays for identification of potent pathogen inhibitors [31,33,51,56,59,60]. In this study, we propose the combination of two HTP antagonistic methods to improve confident selection of potent biocontrol bacterial strains. While the antagonistic co-cultivation method accounts for the entire repertoire of direct inhibitory mechanisms displayed by a biocontrol strain, such as bioactive compounds and competition for nutrients and space, the antagonistic supernatant method pinpoints the inhibitory effect of secreted metabolites such as enzymes, lipopeptides and polyketides. Thus, the two methods provide different results that in combination may aid further mechanistic elucidation. In addition, while the antagonistic co-inoculation method may facilitate the identification of potent candidates ideal for in furrow application or seed coating, where pathogens and biocontrol agents actively compete, the antagonistic supernatant method identifies high producers of bioactive metabolites, which would be advantageous to develop a liquid formulation for foliar product application. In light of the results obtained and considering the different mechanistic aspects involved in pathogenic inhibition, we argue that the combination of our novel antagonistic co-cultivation and supernatant methods constitutes an improved strategy for biocontrol strain identification. The combination of the two presented methods: 1) confidently reflects the fungal inhibition capacity of biocontrol candidates, 2) facilitates HTP screening of large strain collections, 3) can provide valuable insights into type of inhibition mechanisms for further studies and, 4) allows easy comparison of strains by accurate quantification of their inhibition potency.

## Materials and Methods

### Microbial species and growth conditions

The fungal and bacterial strains used in the inhibition assays are shown in Table 1. Fungal species were cultivated on potato dextrose agar (PDA) medium ([Carl Roth]; potato infusion 4 g/l, glucose 20 g/l, agar 15 g/l, pH value 5.6 ± 0.2). For inhibition assays on PDA, cultures were incubated at room temperature with natural light. For inhibition assays in broth, fungal spores were inoculated in potato dextrose broth (PDB) medium ([Carl Roth]; potato infusion 6.5 g/l, glucose 20 g/l, pH value 5.6 ± 0.2) and incubated at 25°C in darkness. *Bacilli* were grown overnight in Lysogeny Broth ([Difco]; Bacto tryptone 10 g/l, [Oxoid]; yeast extract 5 g/l, [Merck]; NaCl 10 g/l, pH value 7.2 ± 0.2) at 37°C with agitation at 250 rpm.

**Table 1:**
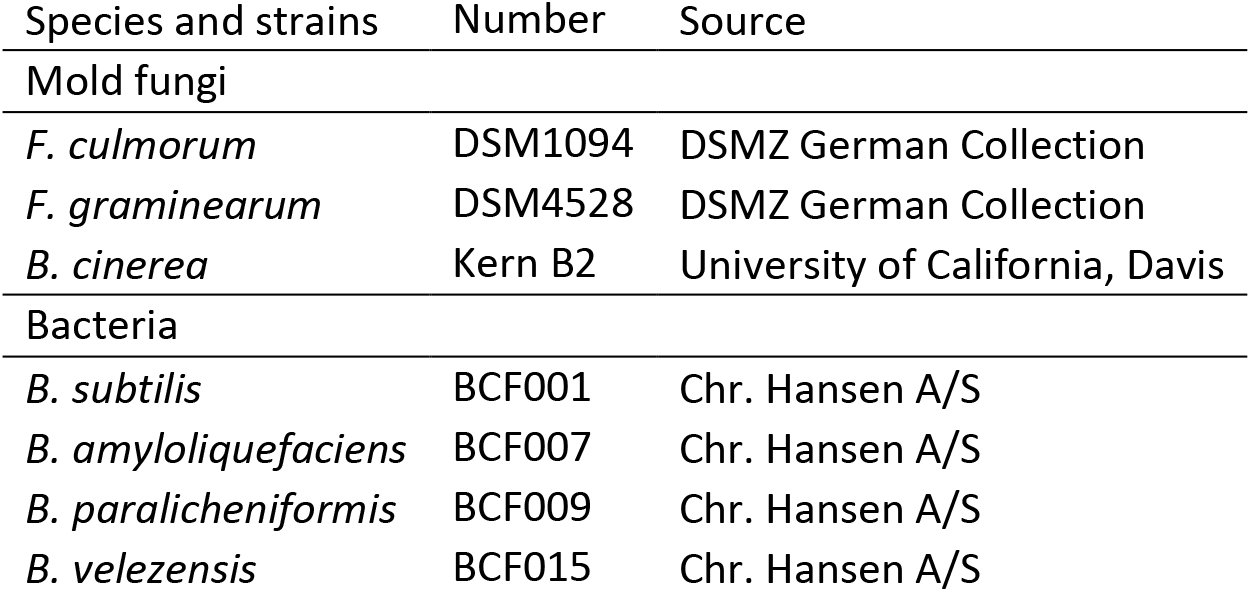
Fungal and bacterial strains used in the present study.

### Fungal spore harvest

Spores were harvested as described by Benoit et al., 2015 and Kjeldgaard et al., 2019. In brief, PDA plates were inoculated with fungi and incubated at 22°C with 16h light/8h dark cycles for at least 2 weeks. Fungal spores were harvested from PDA plates using saline tween solution (8 g/l NaCl, 0.05 ml/l Tween 80) and gentle scraping with an L-spreader. The spore solution was filtered through double layered Miracloth [Millipore] and pelleted by centrifugation at 5000 rpm for 10 min. The supernatant was discarded and the spore pellet resuspended in saline tween. Spore stocks were prepared by adding an appropriate concentration of glycerol.

### Quantification of fungal growth inhibition by co-inoculation with bacterial dilution series

The fungal inhibition assay was prepared in 48-microtiter plates with 0.5 ml PDA in each well. *Bacillus* cultures were adjusted to 2*10^−2^ or 8*10^−4^ at OD_600_ and 5-fold dilution series were prepared with 6 steps using peptone saline as diluent ([Millipore]; Maximum Recovery Diluent 9.5 g/L). 15 μl of each bacterial dilution was inoculated in consecutive wells by spotting the solution in the center of the wells. One strain was assigned per column. Following, a fungal spore solution was prepared with peptone saline and vortexed vigorously to disperse clumps of spores. 15 μl fungal spore solution was co-inoculated with the bacterial dilutions in each well. The approximate final fungal spore concentration of *F. culmorum, F. graminearum* and *B. cinerea* was 5.5*10^4^ spores/ml, 3.1*10^3^ spores/ml and 7.5*10^5^ spores/ml, respectively. Co-inoculated plates were sealed with 3M tape ([Millipore] 0.5 cm) to reduce growth differences between inner and peripheric wells, and incubated at room temperature under natural light conditions. After 5 days, the fungal inhibition was evaluated first by visual inspection and later scored by scanning the plates. The minimal inhibitory cell concentration (MICC) was identified for each strain and averaged between technical duplicates, then averaged between biological triplicates. The MICC (CFU) against each fungal species was converted to an inhibition score by Equation #1 and statistical significance was calculated by two-tailed student’s T-test.

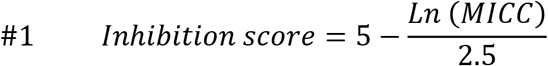

### Quantification of fungal inhibition potency by metabolites in bacterial (cell-inactive) supernatants

The fungal inhibition assay was prepared in 48-microtiter plates with 0.5 ml PDB medium. Spores of *F. culmorum*, *F. graminearum* or *B. cinerea* were added to a final concentration of 1.1*10^4^ spores/ml, 6.1*10^2^ spores/ml and 7.5*10^4^ spores/ml respectively. The spore suspensions were mixed thoroughly to ensure a homogeneous distribution. *Bacillus* spp. cultures were adjusted to OD_600_ of 2 and centrifuged to pellet the cells. The supernatants were collected and the cell pellets discarded. Increasing bacterial supernatant volumes (ranging from 10-80 μl) were inoculated into the fungal spore suspension. Two approaches were implemented for sterilization of the supernatant to omit bacterial growth. Either the bacteriostatic antibiotics chloramphenicol and tetracycline were added to the fungal spore suspension to a final concentration of 50 μg/ml and 10 μg/ml, respectively, or the supernatants were sterilized by filtration ([Sartorious] Minisart Syringe Filter 0.2 μm) prior to co-inoculation with the fungal spores. The plates were sealed with 3M tape and incubated at 25°C without shaking in the darkness. The fungal growth was quantified by spectrophotometric measurements (OD_600_) after 48h for *F. graminearum* and *B. cinerea* inhibition assays and after 67-72h for *F. culmorum* inhibition assays. For more accurate fungal growth estimation, the OD_600_ was measured using a 5×5 well-scanning matrix. Following the evaluation of fungal growth, the supernatants were collected and plated on LB with fungicides (50 mg/L nystatin) to check for unwanted bacterial growth. Statistical analyses were done to compare to the effect of 1) bacterial culture supernatants on fungal growth, 2) filtration and antibiotics on the bacterial supernatants potency, and 3) prokaryotic antibiotics on fungal growth. Statistical significance was calculated by students T-test (two-tailed). A numerical inhibition score was assigned to each strain by quantifying the fungal growth in response to supernatant addition (5 volumes). The inhibition scores were calculated by subtracting the accumulated relative fungal growth from total potential fungal growth (5 conditions tested) as shown in Equation #2:

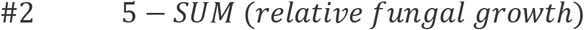

## Supplementary

**Table S1:** The minimal inhibitory cell concentration (MICC) was determined for *B. subtilis* BCF001, *B. amyloliquefaciens* BCF007, *B. velezensis* BCF015 and *B. paralicheniformis* BCF009, as the lower number of cells (CFUs) that inhibit growth of *F. culmorum, F. graminearum* and *B. cinerea* by co-cultivation. The MICC against each fungal species was converted to an inhibition score by formula #1. For quantification of the culture supernatant inhibition potency, an inhibition score was calculated by formula #2 for each bacterial strain and for each experimental cell-inactivation method.

|  | BCF001 | BCF007 | BCF009 | BCF015 |
| --- | --- | --- | --- | --- |
| MICC (CFU) |  |  |  |  |
| *F. culmorum* | 19 | <3 | >3.89*10<sup>4</sup> | <5 |
| *F. graminearum* | 57 | 60 | >1.09*10<sup>5</sup> | 74 |
| *B. cinerea* | 67 | 150 | >1.10*10<sup>5</sup> | 153 |
| Co-cultivation inhibition score |  |  |  |  |
| *F. culmorum* | 3.8 | 4.5 | 0.8 | 4.4 |
| *F. graminearum* | 3.4 | 3.4 | 0.4 | 3.3 |
| *B. cinerea* | 3.3 | 3.0 | 0.4 | 3.0 |
| Supernatant inhibition score (antibiotics) |  |  |  |  |
| *F. culmorum* | 2.63 | 2.81 | 0.62 | 2.06 |
| *F. graminearum* | 2.14 | 1.57 | -0.43 | 1.81 |
| *B. cinerea* | 3.53 | 3.43 | 1.75 | 3.50 |
| Supernatant inhibition score (filtration) |  |  |  |  |
| *F. culmorum* | 2.52 | 2.64 | 0.46 | 2.13 |
| *F. graminearum* | 2.04 | 1.22 | -0.50 | 1.81 |
| *B. cinerea* | 3.55 | 3.42 | 1.32 | 3.53 |
| Supernatant inhibition score (filtration and antibiotics) |  |  |  |  |
| *F. culmorum* | 2.47 | 2.56 | 0.26 | 1.81 |
| *F. graminearum* | 2.35 | 1.37 | -0.40 | 1.72 |
| *B. cinerea* | 3.50 | 3.35 | 0.81 | 3.50 |

**Fig. S1:**
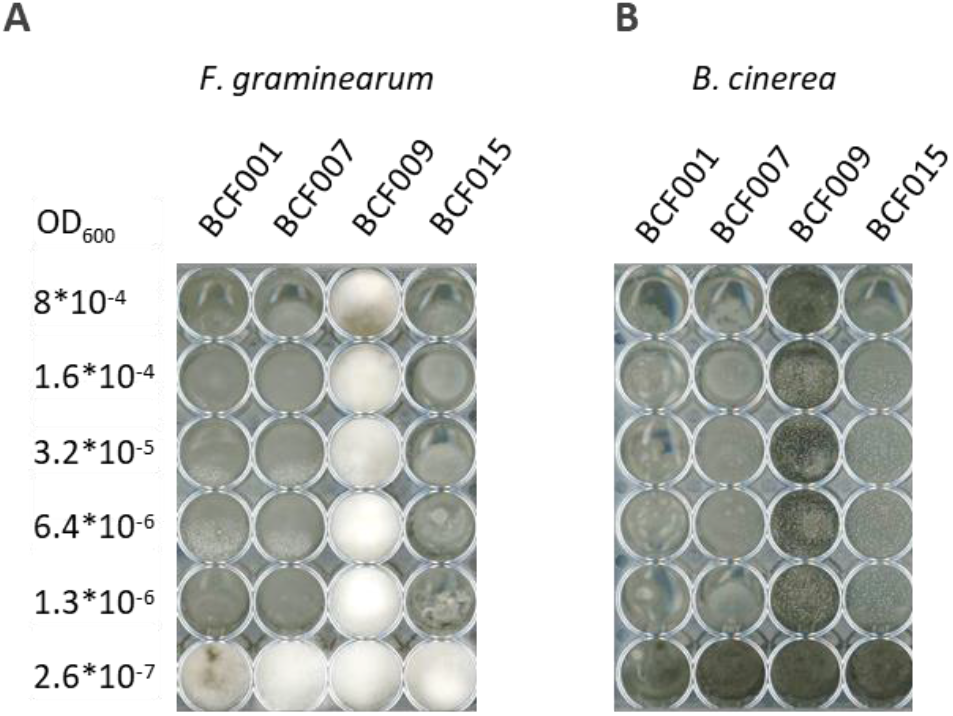
Fungal inhibition assay with *Bacillus* dilution series. Dilution series of *B. subtilis* BCF001, *B. amyloliquefaciens* BCF007, *B. paralicheniformis* BCF009 and *B. velezensis* BCF015 were prepared and inoculated in consecutive columns of a 48-well microtiter plate. In each well, a constant spore concentration was inoculated of (A) *F. graminearum* and (B) *B.* cinerea. Pictures are representative of triplicates.

**Fig. S2:**
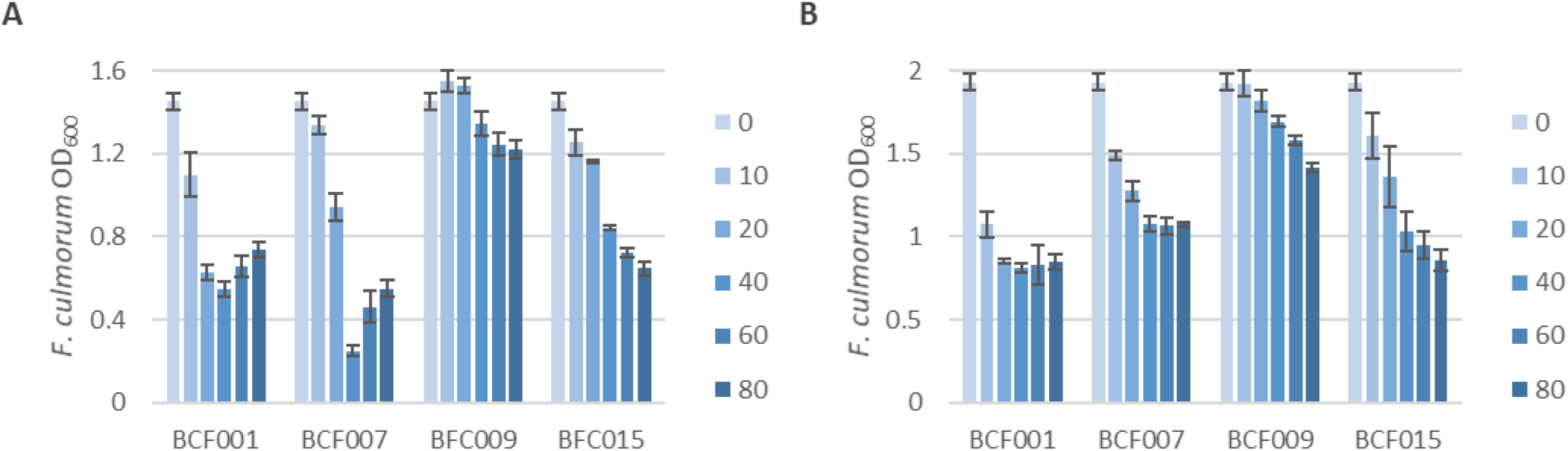
Variation of *F. culmorum* growth with *Bacillus* supernatants. Inhibition assays with *F. culmorum* were prepared from different spore solutions and reached different final growth after 67h (A) and 72h (B) cultivation in PDB medium both with and without bacterial culture supernatants. Growth was evaluated by spectrophotometric measurements (OD_600_). Standard deviations were calculated from biological triplicates.

**Fig. S3:**
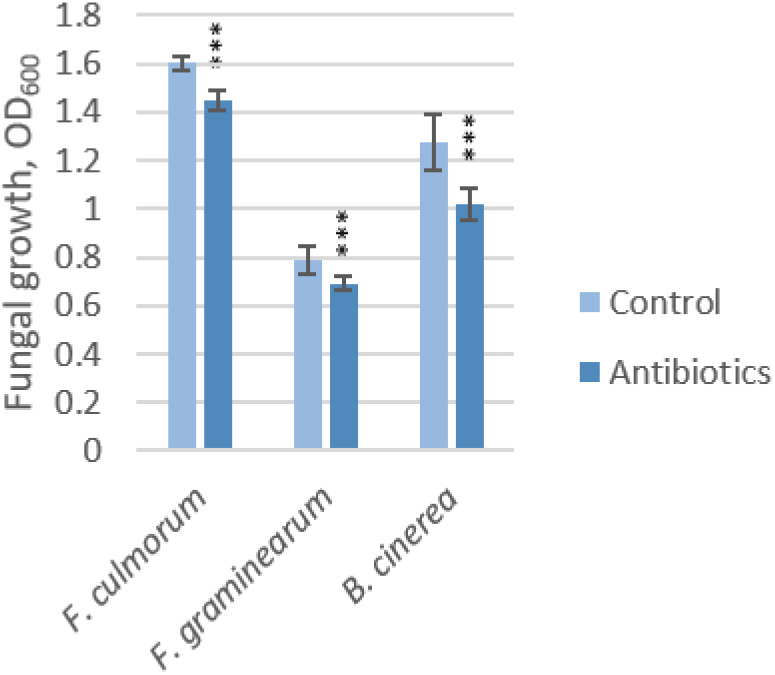
Growth of *F. culmorum, F. graminearum* and *B. cinerea* without and without addition of antibiotics (50 μg/ml chloramphenicol and 10 μg/ml tetracycline). ***P<0.0005

**Table S2:** Statistical significance of different *Bacillus* supernatant sterilization methods on fungal inhibition results. Statistical significance was calculated using student’s t-test. Statistical significant values (p<0.05) are indicated in red.

| Filtered supernatants with antibiotics compared to supernatants with antibiotics |  |  |  |  |  |  |
| --- | --- | --- | --- | --- | --- | --- |
| Supernatant volume (µl) |  | 10 | 20 | 40 | 60 | 80 |
| *F. culmorum* | BCF001 | 0.43 | 0.55 | 0.69 | 0.18 | 0.12 |
|  | BCF007 | 0.21 | 0.01 | 0.24 | 0.82 | 0.11 |
|  | BFC009 | 0.97 | 0.2 | 0.05 | 0.08 | 0.01 |
|  | BFC015 | 0.77 | 0.31 | 0.09 | 0.01 | 0.04 |
| *F. graminearum* | BCF001 | 0.88 | 0.09 | 0.01 | 0 | 0.94 |
|  | BCF007 | 0.31 | 0.14 | 0.66 | 0.85 | 0.06 |
|  | BFC009 | 0.3 | 0.44 | 0.45 | 0.61 | 0.78 |
|  | BFC015 | 0.28 | 0.01 | 0.6 | 0.34 | 0.32 |
| *B. cinerea* | BCF001 | 0.11 | 0.61 | 0.4 | 0.89 | 0.71 |
|  | BCF007 | 0.81 | 0.51 | 0.08 | 0.34 | 0.34 |
|  | BFC009 | 0.07 | 0.1 | 0.02 | 0.1 | 0.22 |
|  | BFC015 | 0.46 | 0.51 | 0.06 | 0.82 | 1 |

Filtered supernatants with antibiotics compared to filtered supernatants
| Supernatant volume (μl) |  | 10 | 20 | 40 | 60 | 80 |
| --- | --- | --- | --- | --- | --- | --- |
| *F. culmorum* | BCF001 | 0.77 | 0.04 | 0.97 | 0.52 | 0.84 |
|  | BCF007 | 0.02 | 0 | 0.06 | 0.02 | 0.87 |
|  | BFC009 | 0.07 | 0.13 | 0.62 | 0.97 | 0.09 |
|  | BFC015 | 0.42 | 0 | 0.18 | 0.09 | 0.04 |
| *F. graminearum* | BCF001 | 0.39 | 0.2 | 0.01 | 0.03 | 0.32 |
|  | BCF007 | 0.42 | 0.75 | 0.08 | 0.05 | 0.98 |
|  | BFC009 | 0.29 | 0.8 | 0.04 | 0.91 | 0.37 |
|  | BFC015 | 0.09 | 0.93 | 0.71 | 0.63 | 0.72 |
| *B. cinerea* | BCF001 | 0.55 | 0.71 | 0.52 | 0.35 | 0.27 |
|  | BCF007 | 0.46 | 0.82 | 0 | 0.11 | 0 |
|  | BFC009 | 0.24 | 0.01 | 0.47 | 0.22 | 0.33 |
|  | BFC015 | 0.15 | 0.89 | 0.03 | 0.35 | 0 |

Filtered supernatants compared to supernatants with antibiotics
| Supernatant volume (μl) |  | 10 | 20 | 40 | 60 | 80 |
| --- | --- | --- | --- | --- | --- | --- |
| *F. culmorum* | BCF001 | 0.38 | 0.29 | 0.7 | 0.04 | 0.16 |
|  | BCF007 | 0.31 | 0 | 0.23 | 0.01 | 0.41 |
|  | BFC009 | 0 | 0.71 | 0.07 | 0.07 | 0.13 |
|  | BFC015 | 0.58 | 0.61 | 0.47 | 0.69 | 0.27 |
| *F. graminearum* | BCF001 | 0.42 | 0.82 | 0.16 | 0.81 | 0.37 |
|  | BCF007 | 0.19 | 0.12 | 0.18 | 0.03 | 0.07 |
|  | BFC009 | 0.19 | 0.54 | 0.14 | 0.42 | 0.24 |
|  | BFC015 | 0.2 | 0.19 | 0.53 | 0.81 | 0.16 |
| *B. cinerea* | BCF001 | 0.2 | 0.32 | 0.35 | 0.5 | 0.44 |
|  | BCF007 | 0.03 | 0.47 | 0.04 | 0.36 | 0 |
|  | BFC009 | 0.29 | 0.47 | 0.06 | 0.13 | 0.56 |
|  | BFC015 | 0.32 | 0.7 | 0.06 | 0.19 | 0.01 |

**Fig. S4:**
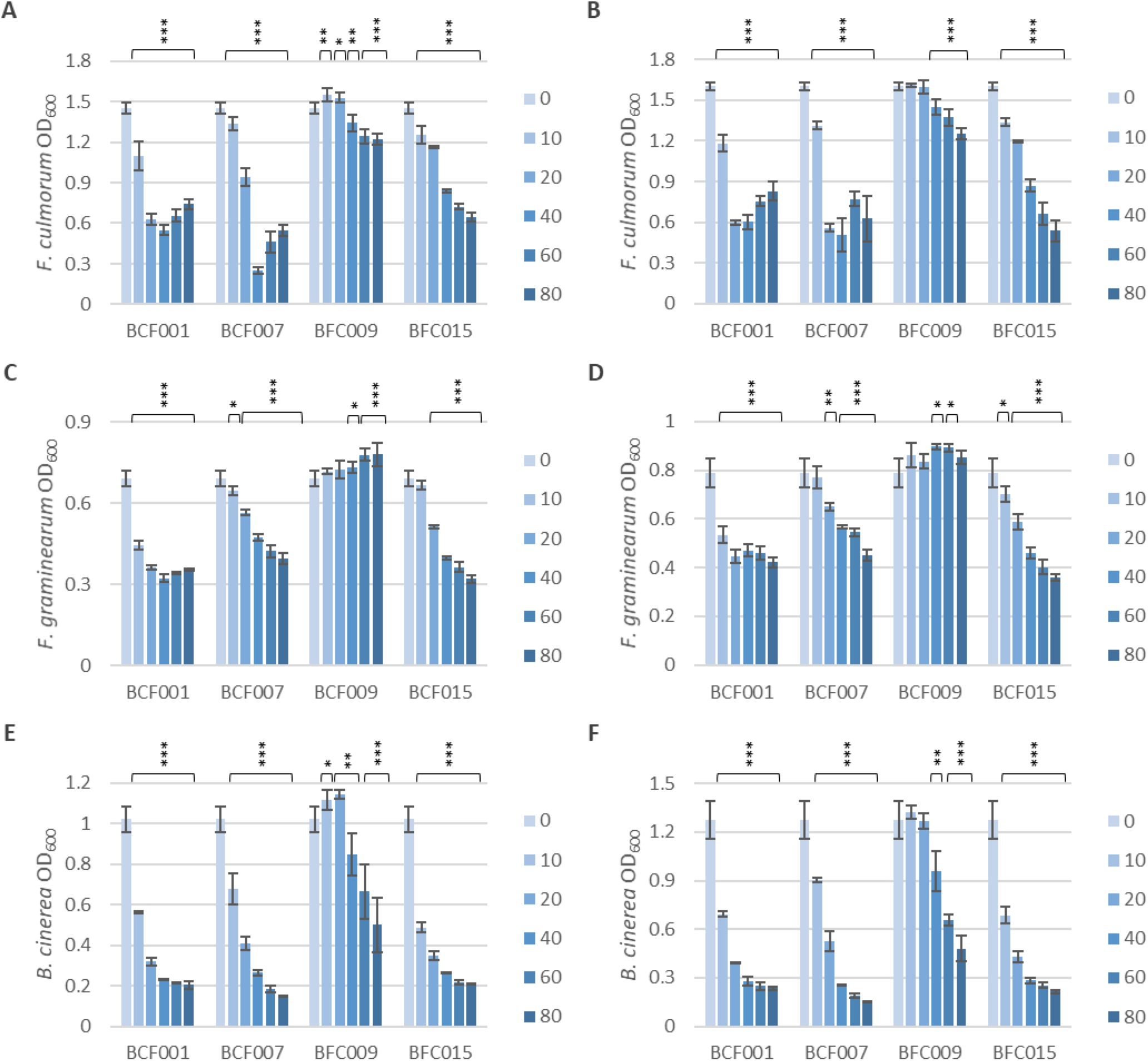
Comparison of methods to circumvent bacterial cell growth in fungal inhibition assays with bacterial supernatants. Fungal spore suspensions of *F. culmorum* (A,B), *F. graminearum* (C,D) and *B. cinerea* (E,F) were inoculated with filtered bacterial culture with antibiotics (A, C, and E) or without addition of antibiotics (B, D, and F). *P<0.05, **P<0.005, ***P<0.0005.

**Fig. S5:**
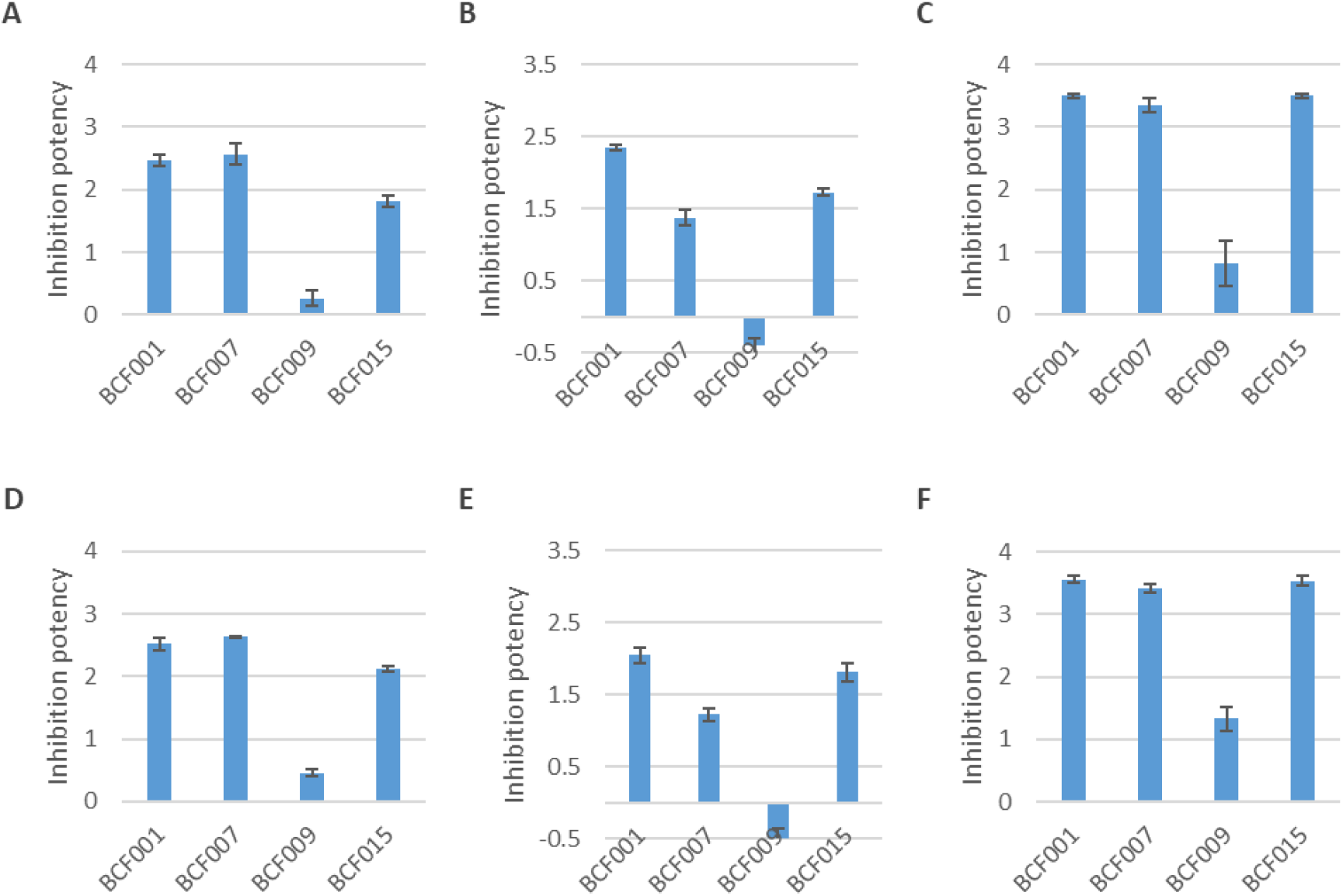
Comparison of supernatant inhibition potencies using *Bacillus* cell inactivation by filtration and antibiotic addition or by filtration only. The culture supernatant inhibition potencies of *B. subtilis* BCF001, *B. amyloliquefaciens* BCF007, *B. paralicheniformis* BCF009 and *B. velezensis* BCF015 against *F. culmorum* (A,D), *F. graminearum* (B,E) and *B. cinerea* (C,F) were calculated by applying the formula #2. The effect of bacterial cell inactivation by filtration and antibiotic addition (A-C) was compared to bacterial cell inactivation by filtration (D-F).

